# Adjusting for principal components of molecular phenotypes induces replicating false positives

**DOI:** 10.1101/120899

**Authors:** Andy Dahl, Vincent Guillemot, Joel Mefford, Hugues Aschard, Noah Zaitlen

## Abstract

High-throughput measurements of molecular phenotypes provide an unprecedented opportunity to model cellular processes and their impact on disease. Such highly-structured data is strongly confounded, and principal components and their variants reliably estimate latent confounders. Conditioning on PCs in downstream analyses is known to improve power and reduce multiple-testing miscalibration and is an indispensable element of thousands of published functional genomic analyses. Further clarifying this approach is of fundamental interest to the genomics and statistics communities. We uncover a novel bias induced by PC conditioning and provide an analytic, deterministic and intuitive approximation. The bias exists because PCs are, roughly, unshielded colliders on a causal path: because PCs partially incorporate a causal genotype effect on one phenotype, the genotype becomes correlated with every phenotype conditional on PCs. We empirically quantify this bias in realistic simulations. For small genetic effects, a nearly negligible bias is observed for all tested PC variants. For large genetic effects, or other differential covariates, dramatic false positives can arise. Though one PC variant (supervised SVA) largely avoids this bias, it is computationally prohibitive genome-wide; further, its immunity to this bias is novel. Our analysis informs best practices for confounder correction in genomic studies.

## 1 Introduction

Association studies of molecular phenotypes have helped uncover genetic regulatory programs underlying a variety of processes including transcription, methylation, chromatin accessibility, translation, ribosomal occupancy, and response to cellular stress. These functional quantitative trait loci (*QTL: eQTL [30, 32], mQTL [33, 29, 7], caQTL [13, 22], pQTL [2], rQTL [8], and reQTL [24, 14]) studies test the associations between genetic variants and molecular phenotypes in a cohort of individuals. For parsimony, we refer only to gene expression phenotypes going forward; however, we emphasize our theoretical arguments and much of our simulations apply to all molecular phenotypes, as well as to other domains with high-dimensional and highly structured data.

These genetic association tests are both performed in local genomic windows (called *cis*) and genome-wide (called *trans*). In *cis*, genetic signals are stronger and simpler, often involving disruption to the physical process of transcription. In *trans*, by contrast, genetic associations are presumably mediated by some complex cellular regulatory process, and so *trans* associations are both biologically central and leave only a subtle and weak signal.

Molecular phenotypes are highly sensitive to structured environmental noise, and this confounding both reduces power and skews the joint distribution of p-values across genes [17, 26]. To partially address these shortcomings, known confounders–such as batch effects–are typically included as covariates in *QTL analyses. But unmeasured confounders–such as subtle experimental variations or cell cycle state–or entirely unexpected sources of confounding often have substantial effect. Fortunately, these confounders intuitively introduce large but low-dimensional variation in the high-dimensional molecular phenotype measurements, and PCs and their variants (which we collectively call CCs, for confounding components) often accurately estimate these unknown factors in practice [26, 36, 31]. As CCs estimate unknown confounders, it is natural to include them as covariates alongside known confounders, and this approach has been shown to decrease power and attenuate joint p-value miscalibration in a wide variety of settings. Domain-specific CCs, like surrogate variables (SVs) [26] or PEER factors [36], have been shown to outperform PCs in eQTL studies, and these methods have become an essential element of thousands of *QTL analysis pipelines [25, 37, 11, 1, 38, 12].

Acknowledging the substantial benefits of CCs, we seek to explore their adverse impact on the false positive rate. The key observation is that CCs are constructed from gene expression measurements that are themselves partially determined by genetic variants and so, inevitably, some causal genetic signal will be captured by CCs. Thus conditioning on CCs is analogous to conditioning on an unshielded collider in a directed graphical model (Figure 1), which generally induces spurious correlations [16, 34]. This unshielded collider bias has become increasingly relevant in modern genetic studies [39, 18, 5, 4]. We show that the bias created by conditioning on PCs results in tests that are misspecified even marginally, are asymptotically inconsistent, and replicate out-of-sample; in contrast, previous discussion of p-value miscalibration focused only on the joint distribution of null p-values, which are not independent in the presence of confounders [26].

**Figure 1:**
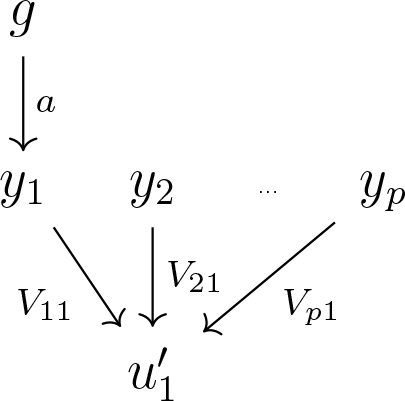
Graphical illustration of the novel bias problem induced by conditioning on CCs. *g* is the genotype affecting one gene expression measurement (*y*_1_). 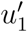 is the first expression PC, which is determined by all *y* and, indirectly, *g*. Conditioning on 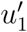 will induce spurious correlation between *g* and all expression measures *y*_*j*_ for *j* ≠ 1.

The PC conditioning bias can be formalized for small genetic effects using standard eigenvector perturbation theory. The approximation to the (suitably defined) bias for testing the *p*-th gene conditional on the first phenotypic PC takes an extremely simple form: 
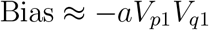
where *a* is the causal effect size, *q* is the gene that is truly causally affected, and *V* are the right singular vectors of the molecular phenotype matrix. Tighter approximations are given below, but the form of this approximation suggests taking the graphical representation in Figure 1 somewhat seriously, as the approximate bias coincides with the bias derived from the graph. A similar, but far smaller, bias can be shown to arise when conditioning on genetic PCs in GWAS, and we suspect similar results may be applicable in other fields.

We then study the practical relevance of this bias using a range of simulations and CC approaches. First, we find that no tested method can correct for this bias even in the extremely simplistic scenario where expression measurements are i.i.d. Gaussian with a small added *cis*-eQTL; this includes PCA, SVA, PEER and linear mixed model approaches [21]. In particular, we show that supervising the CC decompositions with the genotype *g* does not eliminate the bias. We theoretically support this claim for a naive approach that incorporates *g* into PCA by performing PCA on regression residuals. In more complex data with strong confounding, this bias may be drowned out by joint p-value miscalibration and can easily be missed.

In a different regime, where a *trans*-eQTL strongly affects many genes, the bias problem can be massive, even relative to the value added by CC methods. We show this using simulated genotypes and real gene expression data from the GEUVADIS consortium [23] to simultaneously obtain realistic phenotype properties and known false/true positive patterns of genetic association. We show that supervised variants of CC methods, primarily used to improve power, largely solve this bias problem. Identical results hold when the *trans*-eQTL is instead any other known covariate of primary interest; in these differential expression studies, large, transcriptome-wide effects are common. One particularly important feature we identify is that the false positive rate is concave in the number of conditioning CCs, which severely undermines the near-ubiquitous practice of choosing this number to maximize the total number of positive tests.

## 2 The bias due to PC conditioning

This section uses a stylized model of a causal genetic effect added to a background matrix of gene expression values. We imagine a causal genotype vector 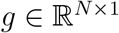 has been measured on *N* individuals and we have measured some baseline gene expression matrix 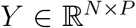 at *P* genes and the same *N* individuals. Our observed gene expression matrix 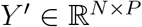 is created by adding a small linear effect of *g* on *Y*: 
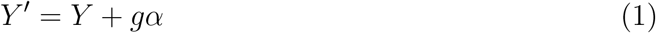
*Y* is considered deterministic but, in this section, fully general, while *g* and the causal effect size 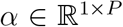 are random. We assume *g* is spherical Gaussian and, for simplicity, that it causally affects only gene *q*. We write this as *α* = *ae*_*q*_, where *a* is the causal effect on gene *q* and *e*_*q*_ is the vector of all zeros except a one in the *q*-th coordinate.

Let 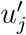 (respectively, *u*_*j*_) be the *j*-th eigenvector of *Y*′ (respectively, *Y*), and let 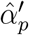 be the regression coefficient on *g* in the regression of 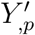 on *g* and 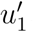, where 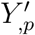 denotes the *p*-th column of *Y*′ (and define 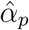 analagously). We aim to assess the impact of conditioning on 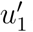 in a regression on *g*. If *u*_1_ were used instead, estimates of *α* will be unbiased as *g* is independent of *Y*.

More specifically, our goal is to quantify the bias induced from *g*’s influence on 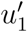: 
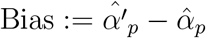

For all *p* ≠ *q*, 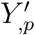 = *Y*_*,p*_, and so the discrepancy between these regressions arises only due to *g*’s effect on 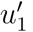.

We assume that *α* is small so that we can use a standard approximation to the perturbed eigenvector (e.g. Section II of [3]): for a generic small perturbation *E*, the first eigenvector of *E* + *Y^T^Y* is approximately 
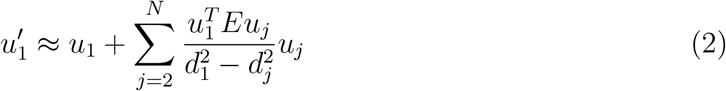
where *d*_*j*_ is the *j*-th singular value of 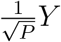. In our case, 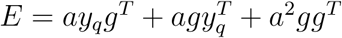, using *y*_*q*_ := *Y*_*,q*_ as shorthand for the *q*-th column of *Y*. Plugging this in to the perturbation approximation gives

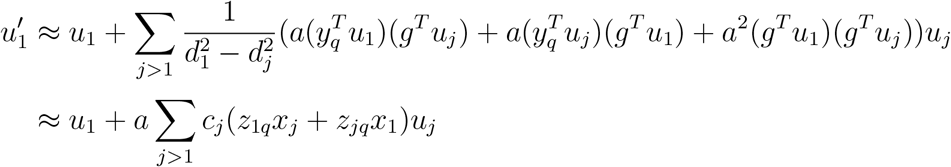

The second line drops *O*(*a*^2^) terms and uses the following simplifying definitions based on the SVD 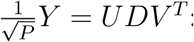:

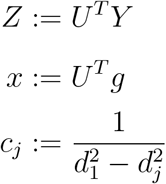

Note that *x* is still a spherical Gaussian random variable.

As an aside, the perturbation approximation is first order in *E* which, itself, depends on *a*^2^. However, a proper second order expansion in *a* would also require incorporating *O*(*a*^2^) terms from a second-order expansion in *E*, hence we drop *O*(*a*^2^) terms from our approximations.

Before turning to the regression estimates, we evaluate a few helpful terms: 
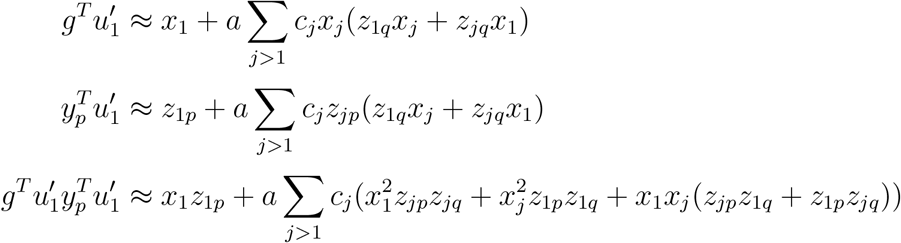

These terms are useful in expanding the bias, which we do using standard two-stage least squares expressions for the regression coefficients: 
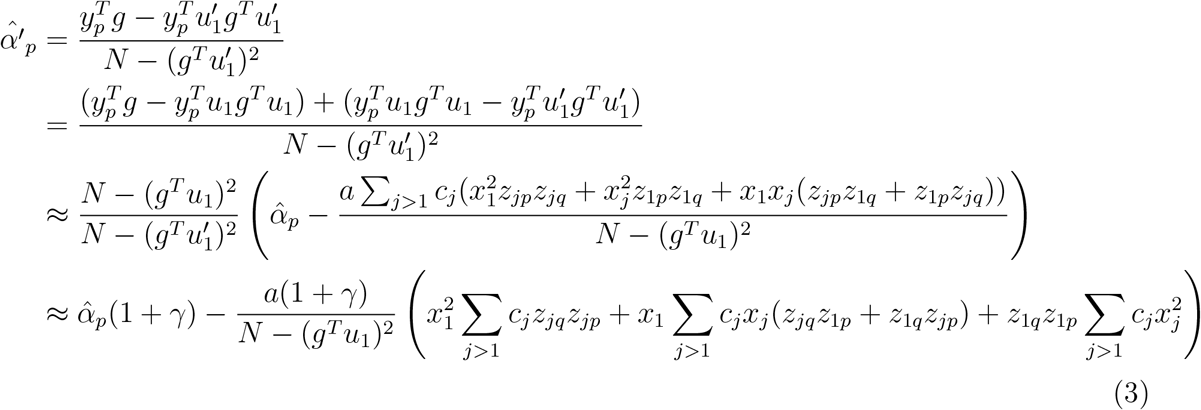
where the approximations are correct to first order in *a*. The last line introduced *γ*, which is defined as 
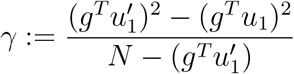

This quantity is written in simple terms and shown to be negligible in the Appendix and ignored going forward.

The bias then becomes 
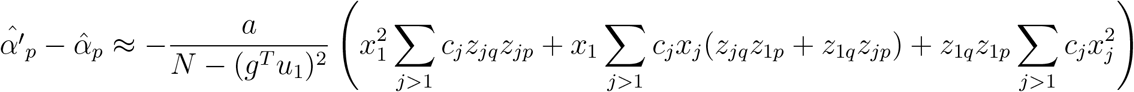

We now drop middle term inside the parentheses that sums terms proportional to *x*_1_*x*_*j*_. These summands are each products of normal random variables with mean zero and only have standard deviation *c*_*j*_(*z*_*jq*_*z*_1*p*_ + *z*_1*q*_*z*_*jp*_). In contrast, the summands in the third term in the parentheses have mean *c*_*j*_*z*_1*p*_*z*_1*q*_ and variance 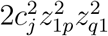. By the central limit theorem, for large *N* the comparison simplifies to comparing a 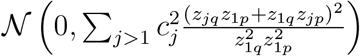 to a 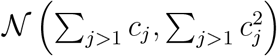. We say the former is negligible because its standard deviation is smaller than the mean of the latter: 
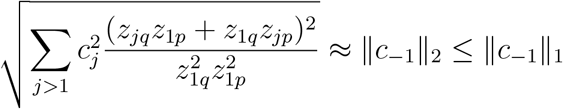

The above approximation assumes 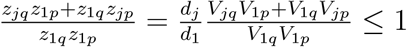, which is reasonable so long as *V* is not too sparse. The inequality is fully general and in our case holds loosely: in the GEUVADIS data (described below) and Marchenko-Pastur spectra with the same aspect ratio, 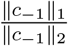 is 19.3 and 9.1, respectively.

Dropping the middle summand simplifies the expression considerably, giving 
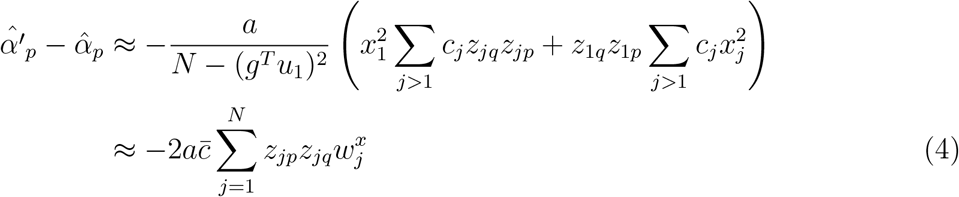

The last line defines the weights *w* on each PC and the overall magnitude term 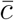 by 
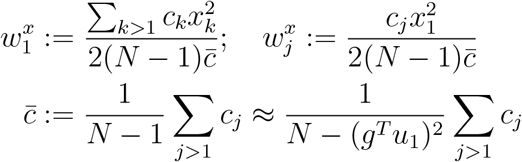

The approximation for 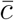 uses 
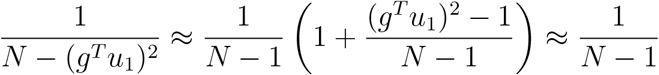
which is correct to first order in the random variable 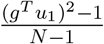, which has mean zero and variance 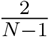.

The *w^x^* partition the perturbation effect among PCs. They are random and depend on the correlation between the random genotype *g* and each PC (i.e. *x*). They are nonnegative and, while they do not sum to one, they do sum to one in expectation. 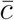, on the other hand, is deterministic and depends only on the spectrum of the baseline expression matrix *Y*. 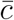 is a condition number and measures the susceptibility of the first eigenvector to perturbations, which is a natural scale factor for the bias induced by perturbing the top PC.

Because of these properties of *w^x^*, the bias in (4) is a weighted correlation between the projections of *p* and *q*–the tested and the causal genes–onto the eigen-axes, i.e. 
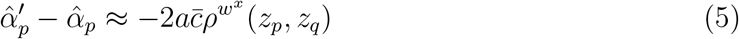
where *ρ^π^* is the correlation between vectors weighting enries by *π*. In particular, *ρ*^1_*N*_^ gives the ordinary correlation between genes *p* and *q* (with or without rotating to eigen-axes). In contrast, *ρ^w^x^^* randomly weights the eigen-axes, but with far greatest weight on component 1 and successively less weight to subsequent axes. The very large weight on component 1 is natural given the regression we consider conditions only on the first component. The successively lower weight on subsequent components is also expected, as perturbation theory argues that eigenvectors with distant eigenvalues are unlikely to interact upon subtle system modifications.

*ρ^w^x^^* is the only remaining randomness in the system because nothing else depends on *g*. This means that, to first order in *a*, choice of *g* affects the bias for testing gene *p* only by defining the relevant notion of similarity between gene *p* and the causally affected gene *q*. In expectation over *g*, 
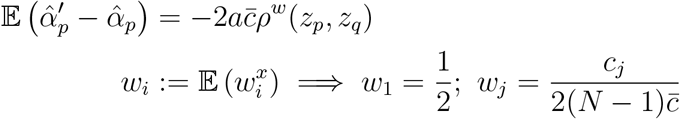

In general, 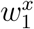 should be very well approximated by its expectation because it is an average over *N* − 1 variables. The other entries of *w^x^*, however, scale with only one (common) random variable, 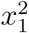; fortunately, this excess variance is counteracted by the fact that the are small (they are collectively as large as 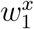). In the GEUVADIS gene expression data (described below), 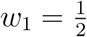, while *w*_2_ = 0.003. In Marchenko-Pastur data with (asymptotic) aspect ratio equal to the GEUVADIS aspect ratio, however, 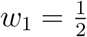 and *w*_2_ = 0.022, confirming that this approximation is better for increasing data confounding.

A final approximation can be made by dropping *w*_*j*_ terms because *w*_1_ ≫ *w*_*j*_ (ignoring the fact that, together, the *w*_*j*_ are as large as *w*_1_): 
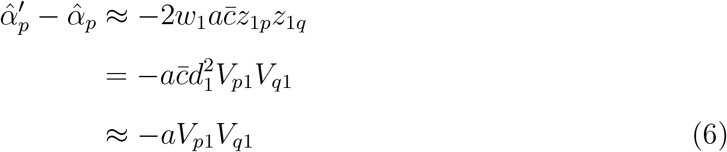

The final line uses the relatively loose approximation that 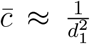, which is derived in equation (7) in the Appendix.

While 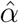 is biased conditional on *α*, it is worth noting that this conditional bias itself has mean zero after averaging over *α*. Equivalently, when *α* = *ae*_*q*_, the bias is zero after averaging over *q* because *V*_,1_ is mean zero For the same reason, the bias is zero on average over *p*. There is no transcriptome-wide average bias or expected bias without knowing the causal gene. Nonetheless, we feel our definition of bias that is conditional on *p* and *q* properly conveys the fact that *p* and *q* are not meaningless indices but biologically determined and replicable out-of-sample.

We have not attempted to generalize the bias calculation to conditioning on more than one PC. However, we suspect an analogous result will hold after modifying *w*, presumably retaining the properties that entries decrease and that the first *K* entries are qualitatively larger than the rest. If this is correct, the weighted correlation will increasingly look like the ordinary correlation as *K* grows. This suggests that the (ordinary) correlation between causal and tested traits is a good intuitive proxy for the extent (and direction) of the regression bias.

### 2.1 Accuracy of the approximations

To empirically assess the quality of the above approximations, we used a prominent and high-quality RNA-sequencing dataset from the GEUVADIS consortium [23]. We first obtained the raw transcript reads from the European individuals in the GEUVADIS consortium. These were then mapped to gene transcripts by aligning the raw reads to the reference hg19 transcriptome using RSEM [28]. We then removed perfectly correlated genes and quantile-normalized genes to standard normal. The resulting matrix has *N* = 375 samples (rows) and *P* = 13, 120 genes (columns). This matrix of gene expression values was used as the baseline expression matrix *Y* both for the simulations in this section and the *trans* simulations described in section 4.

Given the baseline *Y* matrix, we simulated 1,000 independent datasets from the model in (1) with 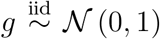 and *α* = *ae*_*q*_, where the causal gene *q* was chosen uniformly at random from {1,…, *P*} and the effect size *a* was varied. For each dataset, the bias 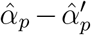 is then computed for each gene *p* ≠ *q*, along with our theoretical approximations to this bias. Finally, the observed biases are regressed on the theoretical biases for each dataset, and we store the regression coefficient and *R*^2^ values.

For both the “full” approximation given in (3) and the simplest approximation given in (6), the median *R*^2^ is greater than .9999 for all *a*. The empirical distribution of these regression coefficients is shown in Figure 2. The median coefficients are always negligibly far from 1, though the empirical 95% confidence intervals are nontrivial. These observations suggest our bias approximation is off by a scale factor near one, possibly because higher-order terms also predominantly scale in *V*_*p*1_*V*_*q*1_. We have not pursued more accurate approximations as the goal is only to demonstrate the existence and qualitative behavior of the bias; moreover, where the perturbation is truly small, we show in simulations below that the resulting bias is essentially negligible.

**Figure 2:**
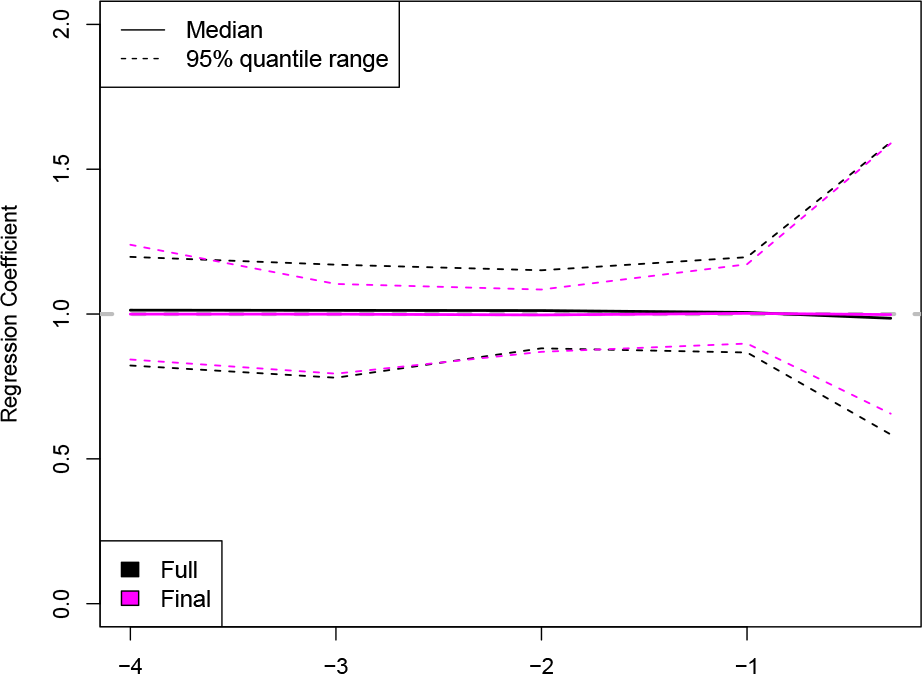
Results from regressing the observed bias in 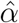 on two novel approximations to the bias. The “final” approximation simplifies several negligible terms in the “full” approximation. The distribution of the regression coefficient is shown, which is identically 1 for perfect approximations.

Overall, the final approximation given in (6) appears to be a very good estimator, despite depending only on *a, q* and *V*_,1_ (when *d*_1_ is large).

### 2.2 Inconsistency and spurious replication

As the approximations developed above depend on the data only through the top right singular vector when *d*_1_ ≫ *d*_2_, datasets with similar *V*_,1_ will have similar regression bias. This is because *a* and *q* are determined by nature.

One implication is that as *N* grows large, the bias stays constant but the regression standard error shrinks, resulting asymptotically in rejecting the null hypothesis transciptomewide for any *g* causally affecting even one gene. This argument implicitly assumes the large-*N* limit for *V*_,1_ (or at least its distribution) is defined and non-sparse (respectively, a.s. non-sparse). Though we avoid formalizing this large-*N* limit, it seems reasonable that if one large confounder is present and that *N* grows large, then *V*_1_, will converge to the confounder and *d*_−1_ will only shrink.

A related implication is that the regression bias will replicate in a new dataset with the same top right singular vectors. There is no requirement on the two sets of left singular vectors to obtain this spurious replication. Again avoiding formality, the top right singular vectors will be similar in datasets with similar confounding structure, and such shared structure is common in molecular phenotypic assays. Further, only sign consistency is assessed in genomic replication analyses, which reduces the burden to prove false positive replication: the datasets need not have equal *V*_*p*1_*V*_*q*1_ terms, only sign-consistency in these terms.

Finally, we note that if the causal signal due to *g* is sufficiently strong, it will dominate leading right singular vectors, which intuitively satisfies the necessary condition for bias replication without any other assumptions on shared confounding across-datasets.

### 2.3 PCA on residuals

An apparent solution to this problem of PC conditioning is to project out *g* before computing PCs. We call this approach supervised PCA (though it is unrelated to [6] and other published definitions). This approach has been studied before through simulations that show supervised SVA provides superior joint p-value recalibration in the presence of confounders [26].

To prove theoretically the flaw in this approach, even for marginal tests when *g* has no causal effect, we first make an assumption: that in realistic data the supervised PCs approximate an approach that projects out *g* from the ordinary PCs, which we call residual PCA. In the GEUVADIS data, described below, we find that the top residual and supervised PC pairs each have squared correlation greater than .99 for independently simulated i.i.d. Gaussian genotypes, suggesting our assumption is reasonable. However, we have not investigated this issue theoretically.

It is easy to prove that conditioning on the residualized PCs spuriously inflates the *t*-statistic for *g*. First, by construction, *g* and the residualized PCs are orthogonal, hence the regression coefficient for *g* is unmodified by conditioning on residualized PCs. The same is true for *g*’s regression coefficient’s standard error, except that the overall regression error 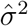 may change. In particular, as PCs (or residual PCs) explain significant variation in *y*_*p*_, 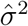 will decrease after conditioning, thus inflating the *t* statistic for *g*’s effect.

The change in *g*’s regression statistics can be directly computed. Let *U* be the first *k* left singular vectors of *Y* and assume columns of *Y* are mean zero and variance 1. We consider the regression equation 
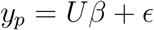

Conditional on only the first *k* PCs, entries of *ε* have mean 0 and variance 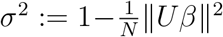; we also assume entries of *ε* are uncorrelated for illustrative purposes. Now let 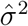 be the ordinary regression estimate of *σ*^2^ conditional on *g* and *U*. If *g* is normalized to length 1, then 
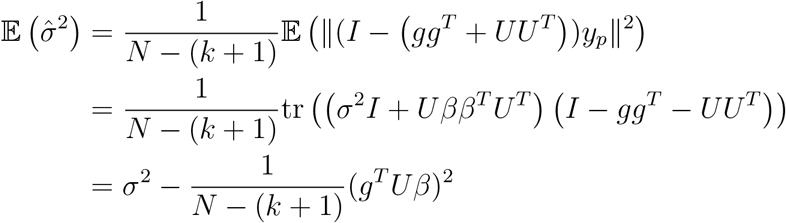

We ignore the second term, which is *O*(*N*^−2^). As *g* is assumed independent of *Y*, the expected value for 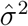 when regressing solely on *g* is just 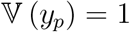. Therefore, conditioning on residual PCs simply inflates *g*’s *t* statistic by roughly a factor of 1/*σ*.

More broadly, this suggests that confounder estimates must strike a delicate balance. Previous sections show how excess correlation with *g* can cause bias, and previous work has shown similar excess correlation leads to loss in power [36]. However, it is now also clear that excess uncorrelation–here, in the form of residualizing *g* from PCs–can lead to different bias problems. We refer to this balance in later sections when discussing the relative performance of different methods, each of which learns different forms of relationships between *g* and estimated confounders.

## 3 *cis*-eQTL simulations with white noise

Having shown that conditioning on expression PCs induces a replicating bias for genetic association tests, we turn now to the practical significance of the problem. We begin with one of the simplest possible simulations based on equation (1): gene expression measurements in *Y* are i.i.d. standard normal; *g* is (indepedently) i.i.d. standard normal; and *α* = *ae*_*q*_, dictating that *g* is a *cis*-eQTL affecting only gene *q*; finally, *q* is drawn, independently of *g* and *Y*, uniformly from {1,…, *P*}, and the effect size *a* is varied. The dimensions of *Y, N* = 375 and *P* = 13, 120, were chosen to match the dimensions of the GEUVADIS data. More realistic simulations are presented in the next section; the focus here is on demonstrating the existence of conditioning bias, which is made dramatically easier by eliminating other sources of null miscalibration.

For each simulated dataset, we perform a 1-sided Kolmogorov-Smirnov (KS) test for deflation in the null regression p-values (for regressing *g* on genes in *Y*′ other than *q*). We also include the (2-sided) KS test for these KS p-values in the plots; this is the “nested” or “double” KS test previously used to detect null p-value miscalibration [26, 27], except we use one 1-sided KS test as the bias we define is expected to decrease null p-values.

After simulating *Y*′ and *g*, we test the association between the genotype and each column of *Y*′–other than the causal gene *q*–with ordinary linear regression conditioned on various CCs. The first set are the PCs of *Y*′; the second set are PEER factors [36]; and the third set are surrogate variables [26]. In this simulation setting, however, SVA declares 0 components significant (through internal permutation tests) and stops–this is arguably ideal behavior. (Also, when no covariate was included, we were unable to implement the “irw” algorithm, which is the suggested default and was used for all of the supervised analyses, discussed below.) We also regress *g* on genes in *Y*′ using linear mixed models that include a random effect with covariance *Y′Y′^T^*, as implemented in ICE [21].

We also consider CCs that are computed with knowledge of the tested genotype *g*. Both SVA and PEER have such supervised options to include a covariate alongside the decomposition. We also compare the supervised PCA approach, discussed in section 2.3, that projects *g* out of *Y*′ before PCs are computed. The primary motivation for supervising these decompositions is to avoid factors that are too correlated with *g*, absorbing the signal and attenuating power [26, 36]. A secondary, and countervailing, goal in supervising these decompositions is to avoid factors that are too uncorrelated with *g*; this balance is described in [26] and in more detail in section 2.3.

Figure 3 presents the quantile-quantile plots for the resulting KS p-values. Unsurprisingly, “None”–including no CCs of any kind–delivers well-calibrated p-values. Also unsurprisngly, the PC approach (for *K* = 1 and *K* = 20) causes noticeable bias for the high-heritability simulation in blue, where the genotype explains 90% of the gene’s variation (equivalently, *a* = 9). However, no statistically significant problem is detected for smaller *a*, suggesting the induced bias from PC conditioning can often be negligible. PEER results are also essentially unperturbed for small *a*, but give highly conservative p-values for large *a*. The linear mixed model (LMM) approach–using either REML or ML (not shown)–yields highly conservative p-values.

**Figure 3:**
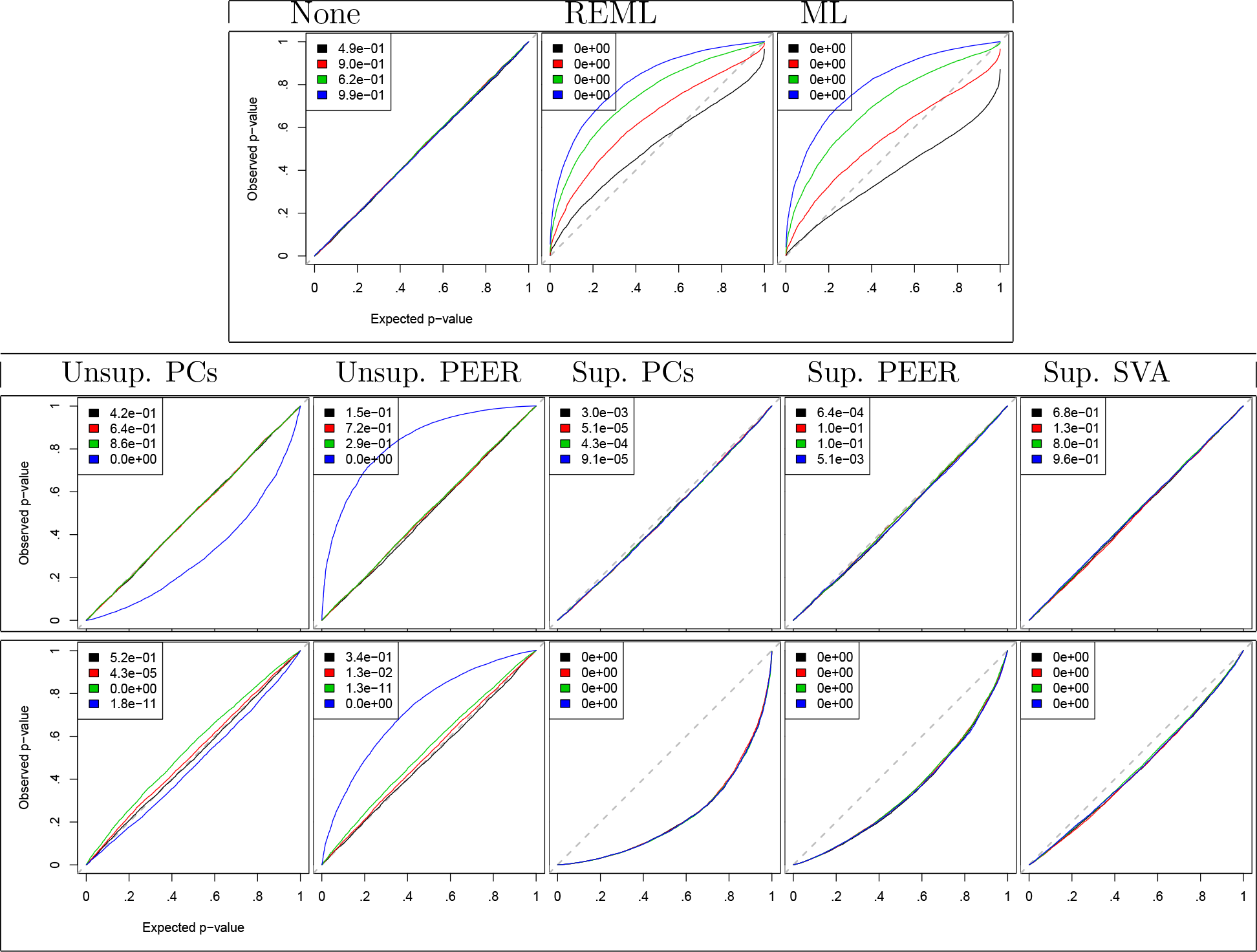
p-value miscalibration for a variety of confounder corrections applied to simulated datasets with a weak genetic effect added to Gaussian white noise. QQ plots for KS tests applied to nominally null p-values are presented, aggregated over 5,000 simulated datasets. KS tests of these KS p-values are presented in the legends. Colors index the strength of the *cis*-eQTL: black corresponds to 0% variation explained by the genotype in the causal gene (the global null); red to 30%; green to 60%; and blue to 90%. The bottom set uses *K* = 1 CC on the top row, *K* = 20 CCs on bottom; the top set uses methods with no *K*.

A different type of bias also appears qualitatively shared among the three supervised methods. The bias grows with *K* (even beyond 20, not shown) and appears independent of the causal effect *a*, persisting even for the global null where *a* = 0. In the case of supervised PCA, this is provably due to supervised PCs being too uncorrelated with the genotype, as noted above. For supervised PEER and SVA, qualitatively similar but attenuated biases are present, suggesting these factors strike a better balance between explaining too much and too little of the signal in *g*. SVA seemed less susceptible to this bias. Some variants of PEER will likely perform better, perhaps including the joint modelling approach in iVBQTL [36] or random effect approach in PANAMA [15]; however, we did not investigate these approaches, which for computational reasons have not been applied to genome-wide scale human data.

Finally, we observed that as *K* grows larger, unsupervised PC adjustment becomes slightly, but significantly, conservative. This effect is only detectable when *a* > 0, i.e. when *g* is truly causal. Surprisingly, the conservativeness it is non-monotonic in *a*: for *K* = 20, 60% PVE (green) gives conservative p-values while 90% (blue) PVE gives anticonservative p-values. This is consistent with some conservative force that emerges for *K >* 1 and scales at a faster rate in *a* than the countervailing anticonservative force we identified theoretically and empirically for *K* = 1. Unsupervised PEER has a related problem, though its conservativeness simply increases in *a*.

Overall, Figure 3 shows that no tested CC method can correct p-value miscalibration in even an extremely simple setting. However, it also shows this problem is very small, especially for small causal effect sizes.

## 4 *trans*-eQTL simulations with real phenotypes

We now present simulations that extend the above *cis*-eQTL simulations in two ways. First, we modified *g* to be a *trans*-eQTL, as *g* now causally explains 30% of the variation in each of 5% of all genes. This means *g* explains 1.5% of variation transcriptome-wide. Null simulations where *g* affects nothing are also included for comparison.

Second, we make the simulation considerably more realistic. We replace the i.i.d. standard normal *Y* matrix by the real GEUVADIS gene expression matrix, described above. Also, as effect sizes are now larger and matching the theoretical bias result is no longer a direct goal, we adopt the standard practice of normalizing columns of *Y*′ to mean zero and variance one. Finally, we simulated *g* to be a binomial SNP genotype with minor allele frequency 20% and then normalized it to mean zero and variance one.

Having established p-value miscalibration in the *cis*-eQTL simulation, we now plot empirical false positive rates in Figure 4. Unlike the KS tests, which assess correlation among gene p-values within-dataset, the FPR assesses a type of mean among the gene p-values. This means the KS miscalibration identified in [26] will only manifest as high variance in the FPR between-datasets. We also show *−* log_10_(*p*)-values averaged over the causally affected genes within each simulated dataset to assess power.

**Figure 4:**
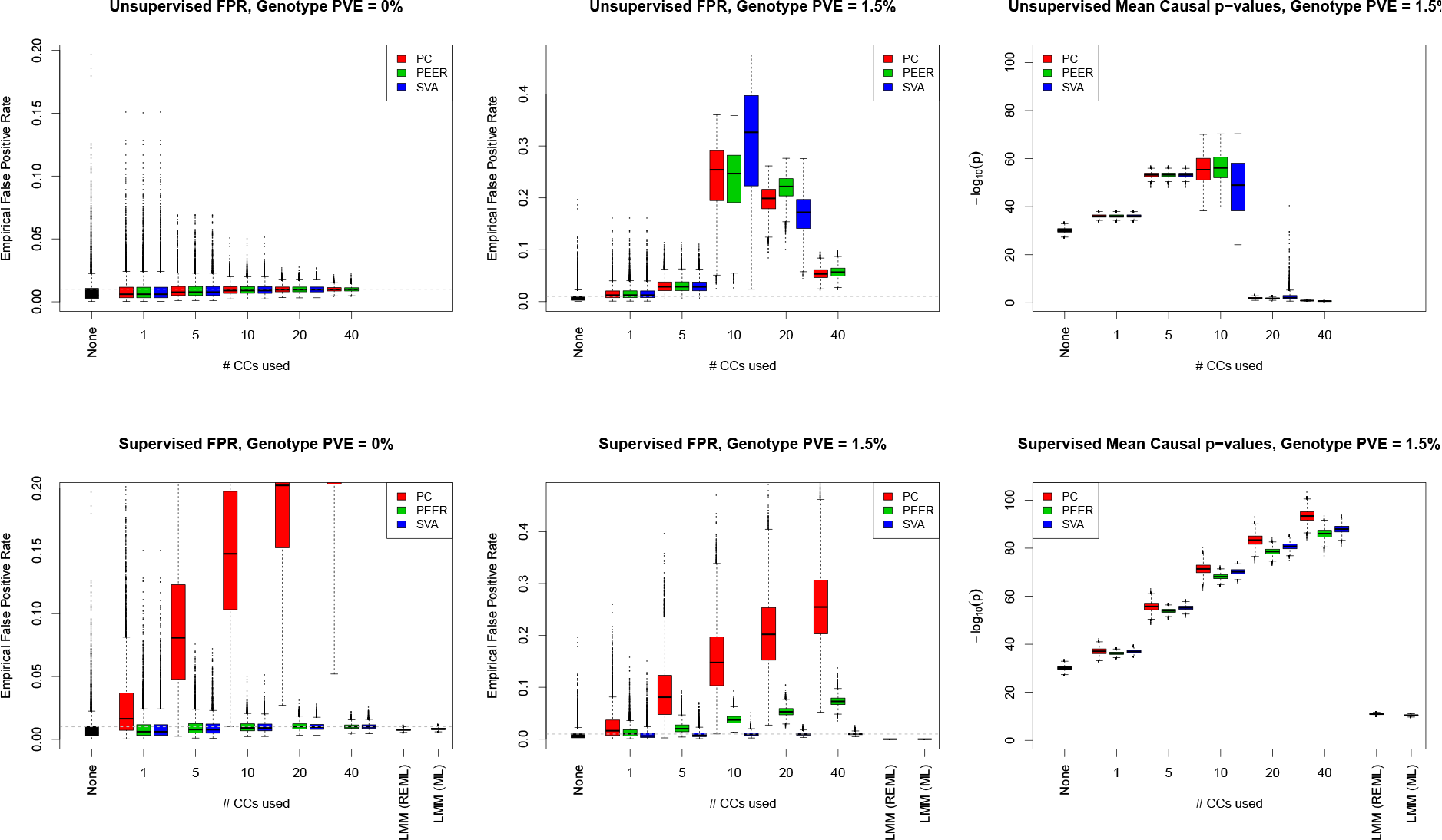
False positive rates and power for a simulated, strong, *trans*-effect added to real gene expression data from GEUVADIS. Results are presented for non-causally affected loci when no (left column) or a large (center column) genetic effect is present; also shown are average − log_10_(*p*)-values for the causally affected genes (right column). Methods that are unsupervised (top) or supervised with respect to the causal genotype (bottom) are both tested. As SVA always deems less than 40 components significant, the corresponding results are absent.

The clearest implication from 4 is that the unsupervised approaches deliver highly inflated false positive rates (top, middle panel) and attenuated power (top, right panel). While the LMM approaches we studied are well-calibrated, they offer even lower power than simply regressing on *g*; however, other LMM approaches may deliver higher power [20]. The obvious conclusion is that one should supervise CCs with all large-effect covariates, roughly yielding well-calibrated FPRs for SVA and slightly inflated FPRs for PEER. However, as is expected from our above theory and is discernible from previous empirical results [26], supervised PCA gives dramatically inflated FPR.

The obvious caveat is that the large-effect covariates must be known in advance. This does not hold when searching for a causal effect among a large set of potential covariates, often outnumbering the sample size, as is standard in genetic studies. While one could refit confounder estimates for each tested *g* in turn, this is typically computationally infeasible; however, pre-screening loci, perhaps by using unsupervised CCs, may dramatically reduce the computational burden. Moreover, practitioners often fail to include known large effect covariates when creating CCs. Finally, to our knowledge, the observation that this supervision eliminates false positives–rather than merely recalibrating the across-gene independence of p-values [26] or increasing power [26, 36]–is novel.

### 4.1 False positive replication

We slightly modified the *trans* simulation to assess the out-of-sample replication rate for false positives. For each simulated dataset, generated exactly as in the above *trans*-simulation, we split the data (*g* and *Y*′) into halves and repeat the above analyses on each half. We compute the replication rate as the fraction of false positive discoveries from the first half that are deemed positive in the second half, using a significance level of *α* = .001 for both splits; we then compute the replication rate after transposing the roles of the two splits. Finally, we average over false replication rates for 5,000 independently simulated datasets and two choices of initial split per dataset (10,000 scenarios in total), discarding scenarios where there are no false discoveries in the initial split.

Results are shown in Table 1 after converting to the replication rate inflation, defined as the ratio of the replication rate to its null expectation (i.e. .001). First, no significant inflation is obtained if confounder corrections are entirely omitted, where significance is defined by the mean inflation being more than 1.65 standard errors greater greater than 1. But both PC and PEER correction substantially inflate the false positive replication rate. For both methods, the using larger *K* exacerbates problem, though supervision improves PEER’s performance and hurts PCA. Unsupervised SVA has inflation problems similar to unsupervised PCA and PEER, but after supervision no statistically significant inflation is detected.

**Table 1:** False positive replication rate inflation after confounder correction. Shown are the false positive replication rates at a significance level of *α* = .001 divided by their expectation. Inflations significantly greater than 1 are indicated with an asterisk.

|  | No CCs | 1 PC | 10 PCs | 1 PEER | 10 PEER | 1 SV | 10 SVs |
| --- | --- | --- | --- | --- | --- | --- | --- |
| Unsupervised | 0.84 | 2.59* | 151.6* | 2.70* | 148.38* | 2.67* | 181.15* |
| Supervised | N/A | 32.47* | 168.69* | 2.02* | 8.22* | 0.68 | 1.09 |

## 5 Discussion

We have shown a novel source of bias induced by conditioning on estimated confounders in association tests. We derived an analytical approximation in the simple case of conditioning on one PC and a small causal effect. We assessed the magnitude of the bias empirically through simulations, with a small effect simulation showing the existence of bias and a large effect simulation demonstrating the impact on the FPR in a realistic setting. This bias will replicate in datasets with similar confounding structures, which is often obtained for high-throughput molecular phenotypes.

This bias problem is related to, but distinct from, known biases caused by confounding. Previous theoretical results have focused on confounding’s impact on the joint null distribution of p-values, showing that an idealized form of correction–i.e. where confounders are perfectly estimated–yields independent p-values for the tests of association between one genotype and each gene [27]. Our results undermine the basic assumptions of this theory, though, by showing that causal effects naturally bias confounder estimates. This novel bias that we show arises from confounder correcting, unlike known biases from uncorrected confounders, results in tests that are misspecified even marginally and that will replicate in similar datasets.

Overall, we find that the supervised versions of SVA and, to a lesser extent, PEER give well-calibrated FPRs and the greatest power among approaches considered. Though no confounder correction method delivers truly independent tests, this effect is generally small and, in the presence of strong confounders, the biases induced by supervised methods are typically negligible compared to the eliminated biases. Further, to avoid computing supervised factors for each SNP genome-wide, *QTL analyses using unsupervised factors can be corrected *post-hoc* by repeating the analysis at significant (or suggestive) genotypes with the appropriately supervised decompositions.

Nonetheless, many high-profile studies fail to supervise their SVA or PEER analyses with respect to strong covariates. This is partially due to a common misconception amongst practitioners that SVA and PEER corrections solely improve power and, thus, that suboptimal implementations are at worst conservative.

There are also more complex scenarios where the appropriate model for covariates is unclear. For example, in gene coexpression studies, covariance or partial covariance can be used to learn complex and subtle graphical models [19, 35]: failing to correct for confounding will yield biologically meaningless confounder networks, while a correction approach with subtle biases may induce a different biologically meaningless network. Latent variable graphical models present a possible solution to this problem [9]. More generally, low-rank-plus-sparse decompositions may be able to simultaneously learn latent confounders and eQTLs transcriptome-wide [10].

We note that qualitatively similar replicating false positive associations can arise when conditioning on unshielded colliders in multiphenotype association tests [4]. This can be expressed in our motivating graphical model in Figure 1 as an environmental node *e* causally affected by several of the phenotype nodes, *y*_*p*_. As *y*_*p*_’s contribution to a generic *e* is arbitrary–unlike its contribution to a PC, which is *O*(*P*^−1/2^)–the replicating false positive problem can be substantially larger than in our context. However, that confounding is entirely dataset-dependent, while our bias is in a sense universal–existing, for example, even when all noise in the *y*_*p*_ is i.i.d. Gaussian. Nonetheless, our false positives will only regularly replicate when the *y*_*p*_ are correlated.

A theoretical question we leave open is whether it is possible to deliver tests that are truly independent across genes in the presence of confounding. The difficulty is clear, as any errors in the estimated confounders will, upon conditioning, induce a fresh source of confounding (albeit one that is potentially better behaved and smaller). Another difficult question is whether the decomposition methods studied can be supervised with respect to a large set of tested covariates, most of which will be entirely null, in a computationally feasible manner. If possible, efficiently updating SVA decompositions for each new added covariate would largely achieve this.

## Appendix: *γ*

Defining and simplifying *γ* gives

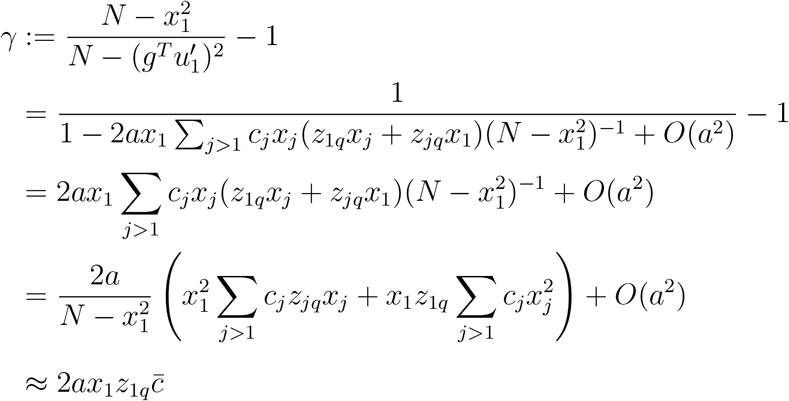

The last line uses the approximations that 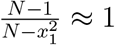 and replaces the sums over *j* by their expectations, which is reasonable as

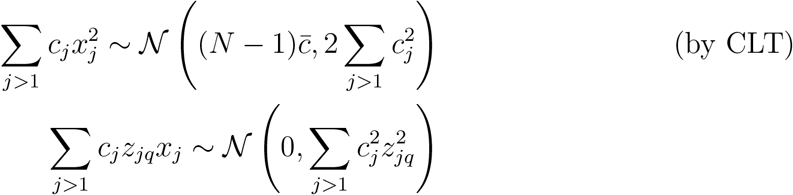

The CLT approximation should be good unless a very small number of eigenvalues (other than the first) are far larger than the rest.

Unfortunately, *K* eigenvalues will be large compared to all other if there are *K* strong confounders, so this approximation will be worse for more realistic data. In Figure 5, for example, the top singular value is roughly 1.5 times larger than the second. Presumably a regression conditioning on *K* PCs would require the last *N − K* to be small, which would be more reasonable.

**Figure 5:**
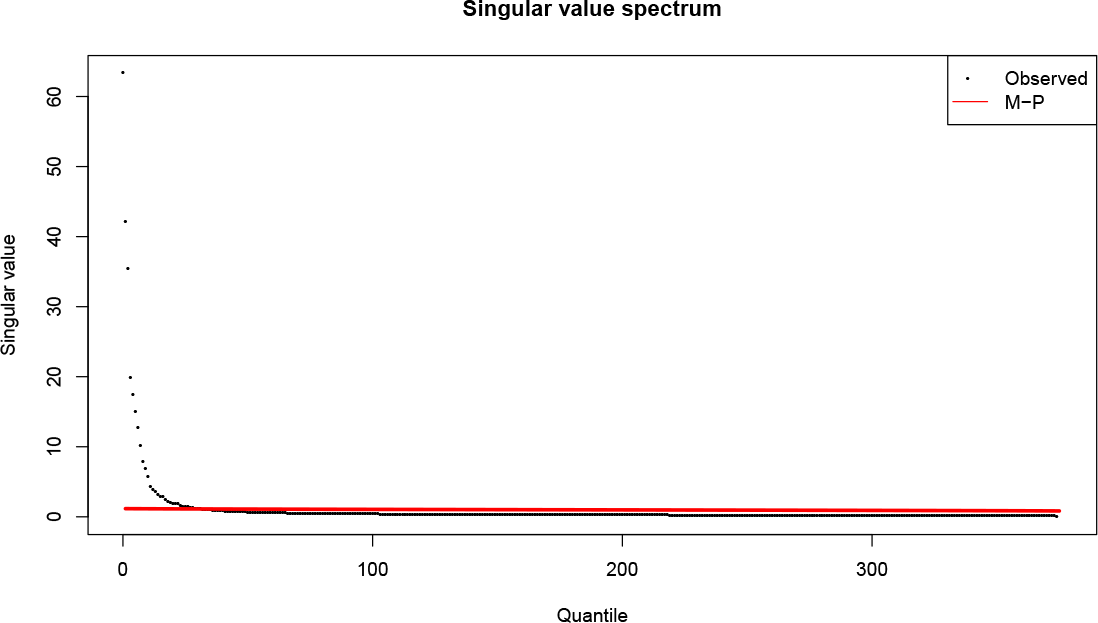
Singular-value spectrum of our process gene expression matrix from GEUVADIS. M-P refers to the Marchenko-Pastur distribution with asymptotic aspect ratio equal matching the GEUVADIS data.

We now derive a loose approximation to 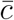 that is also used in the main text:

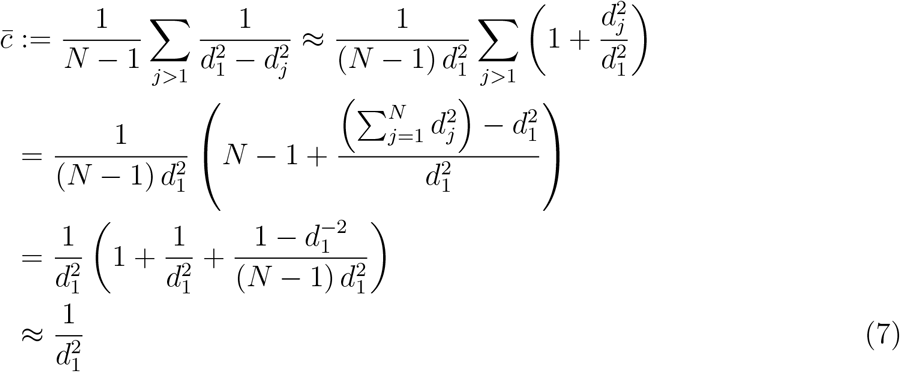

This used the fact that 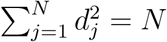, which holds because columns of *Y* have been centered and scaled:

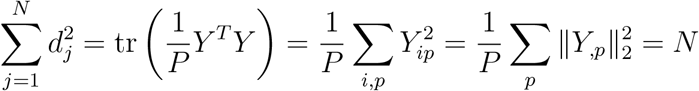

Using (7) to simplify 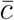, *γ* can then be approximated by 
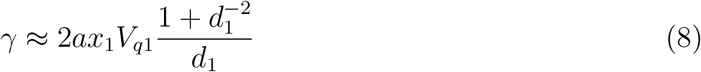

In practice, this term is negligible: *a* is assumed small, *V*_*q*1_ is on the order of *P*^−1/2^, and *d*_1_ tends to be large in real data, e.g. 8 in GEUVADIS.

